# Copper-Free Click Chemistry Enables High-Fidelity Engineering of Mitochondria-Targeted Brain-Derived Exosomes

**DOI:** 10.1101/2025.11.19.689327

**Authors:** Xin Yan, Xinqian Chen, Shanshan Hou, Xiaoxu Li, Jingyue Ju, Zhiying Shan, Lanrong Bi

**Affiliations:** Department of Chemistry, Michigan Technological University, Houghton, Michigan 49931, United States; Health Research Institute, Michigan Technological University, Houghton, Michigan 49931, United States; Department of Kinesiology and Integrative Physiology, Michigan Technological University, Houghton, Michigan 49931, United States; Department of Molecular Pharmacology and Therapeutics and Department of Chemical Engineering, Columbia University, New York City, NY 10027, United States

**Keywords:** Mitochondrial-targeting exosome, SPAAC, CuAAC, Click chemistry, Exosome-based drug delivery

## Abstract

Mitochondrial dysfunction is a hallmark of neurodegenerative and neuroinflammatory disorders, including hypertension and cardiovascular disease, yet strategies for safe and precise mitochondrial-targeted delivery remain limited. Here, we establish strain-promoted azide–alkyne cycloaddition (SPAAC) as a biocompatible, high-fidelity chemical platform for engineering mitochondria-targeted brain-derived exosomes (BR-EVs). Copper-free click conjugation of a mitochondrial-targeting ligand (e.g. Cy5-DBCO) under mild aqueous conditions preserved vesicle morphology (30–150 nm core; 120–200 nm hydrodynamic), proteomic composition, and uptake dynamics. Time-course imaging and fluorescence recovery after photobleaching (FRAP) revealed unaltered endocytic kinetics, >75 % mitochondrial colocalization, and intact organelle architecture. In vivo neuroinflammation and biodistribution analyses demonstrated immunological neutrality, strong central nervous system retention, and minimal peripheral dispersion following intracerebroventricular administration. Proteomic profiling of unlabeled Sprague–Dawley (SD) and hypertensive Dahl salt-sensitive (DSS) BR-EVs uncovered hypertension-driven enrichment of oxidative and complement pathways correlating with mitochondrial fragmentation and reactive oxygen species generation in neuronal cultures. These findings establish SPAAC-mediated ligand conjugation as a biocompatible and chemically precise approach for generating mitochondria-targeted exosomes that preserve exosomal identity, biodistribution, and signaling fidelity—advancing a foundational platform for organelle-specific delivery and mechanistic imaging in the central nervous system.

## Introduction

Mitochondrial dysfunction represents a pivotal pathological mechanism in numerous neurodegenerative diseases, including Alzheimer’s disease (AD), Parkinson’s disease (PD), and amyotrophic lateral sclerosis (ALS).^1–5^ This dysfunction manifests as impaired energy production, excessive generation of reactive oxygen species (ROS), and dysregulated calcium homeostasis, culminating in neuronal loss and progressive cognitive or motor decline. Neurons, with their high metabolic demands, are especially susceptible to these deficits, which impair synaptic plasticity, axonal integrity, and cellular resilience.^1–6^

The mitochondrial aberrations exhibit disease-specific nuances yet converge on shared destructive pathways. In AD, amyloid-β (Aβ) oligomers and hyperphosphorylated tau accumulate at mitochondrial membranes, promoting dynamin-related protein 1 (Drp1)-driven fission. This hyperfission yields fragmented, dysfunctional mitochondria and represses nuclear oxidative phosphorylation (OXPHOS) genes, slashing ATP levels by as much as 50% in hippocampal neurons. ^2,7^ In PD, α-synuclein aggregates inhibit complex I of the ETC, precipitating respiratory failure, ATP exhaustion, and ROS-mediated demise of dopaminergic neurons in the substantia nigra.^8–10^ In ALS, mutations in superoxide dismutase 1 (SOD1) or TDP-43 hinder mitochondrial transport and calcium uptake, eliciting motor neuron hyperexcitability, axonal dystrophy, and relentless paralysis.^11–13^ Across these conditions, the resultant oxidative stress, inflammation, and glial reactivity position mitochondria as an indispensable therapeutic target.

Despite this imperative, mitochondrial-targeted interventions are hindered by two critical barriers: the blood-brain barrier (BBB), which restricts CNS access, and the challenge of achieving organelle-specific delivery within neurons.^14–19^ Conventional carriers like liposomes or polymeric nanoparticles suffer from suboptimal BBB traversal, biocompatibility issues, and off-target effects, while viral vectors raise concerns over immunogenicity and genotoxicity.^18,19^ Exosomes—endogenous nanovesicles (30–150 nm) secreted by diverse cells—emerge as a superior paradigm. These lipid-bilayered structures boast inherent stability, negligible immunogenicity, and natural affinity for neurons and glia, facilitating BBB penetration through receptor-mediated transcytosis.^20–22^ Moreover, their lipid envelopes shield therapeutic payloads (e.g., miRNAs, peptides, or small molecules) from degradation, rendering exosomes a cornerstone for targeted nanomedicine.^20–22^

Yet, unmodified (“naïve”) exosomes display nonspecific biodistribution and superficial cellular engagement, curtailing their potency against mitochondrion-centric pathologies. This underscores the need to engineer mitochondrial-targeting exosomes: by directing payloads precisely to impaired mitochondria, these modified vesicles can amplify therapeutic impact, mitigate off-target toxicity, and restore bioenergetics at the disease epicenter. To confer such precision, surface functionalization with mitochondria-targeting ligands (MTLs)—such as lipophilic cations or peptide motifs—is essential.^23–26^ These ligands exploit the negative membrane potential of mitochondria to drive selective accumulation, promote endosomal escape, and ensure prolonged cargo retention within the organelle. Consequently, MTL-engineered exosomes represent a transformative drug delivery platform,^27^ enabling subcellular resolution in therapies that traditional systems cannot achieve. Their biocompatibility, coupled with scalable production from autologous sources, positions them as a clinically viable avenue to combat neurodegenerative progression

Building on this foundation, our prior work introduced a copper-catalyzed azide-alkyne cycloaddition (CuAAC) method to conjugate a MTL (e.g. a rhodamine derivative) onto BR-EVs, yielding robust mitochondrial colocalization *in vitro* and *in vivo* across cortical and hippocampal regions—without provoking inflammation. ^27^ However, the copper catalyst’s propensity to catalyze Fenton-like reactions and generate hydroxyl radicals poses a translational roadblock, exacerbating oxidative stress in redox-vulnerable neuronal milieus like those in AD and PD.

To surmount this, we herein leverage SPAAC click chemistry—a metal-free, bioorthogonal ligation that sustains conjugation fidelity under mild physiological conditions. By tethering MTLs (e.g., Cy5-DBCO derivatives) to azide-modified BR-EVs via dibenzocyclooctyne (DBCO), SPAAC preserves exosomal structure, cargo viability, and bioactivity while eliminating catalyst-induced cytotoxicity. This innovation not only refines mitochondrial targeting but also bolsters the platform’s safety profile for chronic CNS applications.

Our present study unveils SPAAC-engineered mitochondrial-targeting exosomes as an advanced, biocompatible vector for neuronal organelle delivery. Through rigorous physicochemical profiling, mitochondrial trafficking assays, and functional rescue evaluations under oxidative duress, we demonstrate superior performance relative to CuAAC and naïve counterparts. This work pioneers a clinically attuned exosome-based strategy to redress mitochondrial failure, heralding new horizons in neurodegenerative therapeutics.

## 2. Results and Discussion

### 2.1. Structural Characterization of BR-EVs

Transmission electron microscopy (TEM) analysis of isolated BR-EVs from both normotensive SD and hypertensive DSS rats revealed a homogeneous population of nanosized vesicles displaying the canonical cup-shaped or saucer-like morphology characteristic of endosome-derived exosomes, with core diameters spanning 30–150 nm. These dimensions align with the established size range for exosomes (typically 30–150 nm), which facilitates their role in intercellular communication by enabling efficient membrane fusion and cargo transfer without eliciting strong immune responses. Complementary DLS measurements in hydrated suspensions further characterized the vesicles, yielding hydrodynamic diameters of 120–200 nm, which incorporate the vesicle core, hydration shell, and any adsorbed solutes under physiological conditions. Notably, no discernible structural differences emerged between DSS- and SD-derived BR-EVs, with equivalent core sizes and hydrodynamic profiles. This invariance suggests that hypertension in the DSS model does not perturb core exosomal biogenesis pathways, such as ESCRT-dependent sorting or lipid raft assembly,^28,29^ confining pathological influences on internal cargo alterations—like enriched pro-inflammatory microRNAs or oxidative enzymes—that could modulate recipient cell responses without altering gross morphology. Such structural conservation is advantageous for comparative studies and therapeutic engineering, as it minimizes variability in uptake kinetics or stability; however, it underscores the need for orthogonal assays (e.g., proteomics or lipidomics) to probe subtler disease-specific modifications that might influence bioactivity, such as membrane fluidity or surface charge.

### 2.2 Native Proteomic Landscape of BR-EVs Prior to Surface Engineering

To define the intrinsic molecular composition of BR-EVs before chemical modification, we performed label-free proteomic profiling of EVs isolated from hypertensive DSS and normotensive SD rats. This baseline dataset establishes the native cargo architecture of BR-EVs and provides a reference point for evaluating subsequent bioorthogonal surface functionalization. Across both groups, we identified ∼760 proteins, including 602 shared proteins, 145 DSS-unique proteins, and 13 SD-unique proteins. The markedly greater number of unique proteins in DSS-EVs reflects hypertension-induced remodeling of vesicular cargo, consistent with heightened oxidative stress, chronic inflammation, and mitochondrial vulnerability in the hypertensive brain. In contrast, SD-EVs were enriched in proteins associated with proteostasis and antioxidant defense, indicating preserved homeostatic signaling under normotensive conditions.

Post-translational modification analysis revealed 42 phosphorylation events; all located within shared proteins such as ion ATPases and myelin-associated enzymes. No unique phosphorylation events or ubiquitination, nitrosylation, or SUMOylation sites were detected under the search conditions used. These results indicate that hypertension predominantly alters protein composition rather than global PTM patterns, maintaining a conserved vesicular signaling backbone while selectively exporting stress-adaptive cargo.

Defining the molecular signature of native BR-EVs prior to any chemical modification provides a rigorous foundation for subsequent SPAAC engineering, and this can enable us evaluation of whether surface conjugation preserves intrinsic vesicle identity and biological characteristics.

**Scheme 1:**
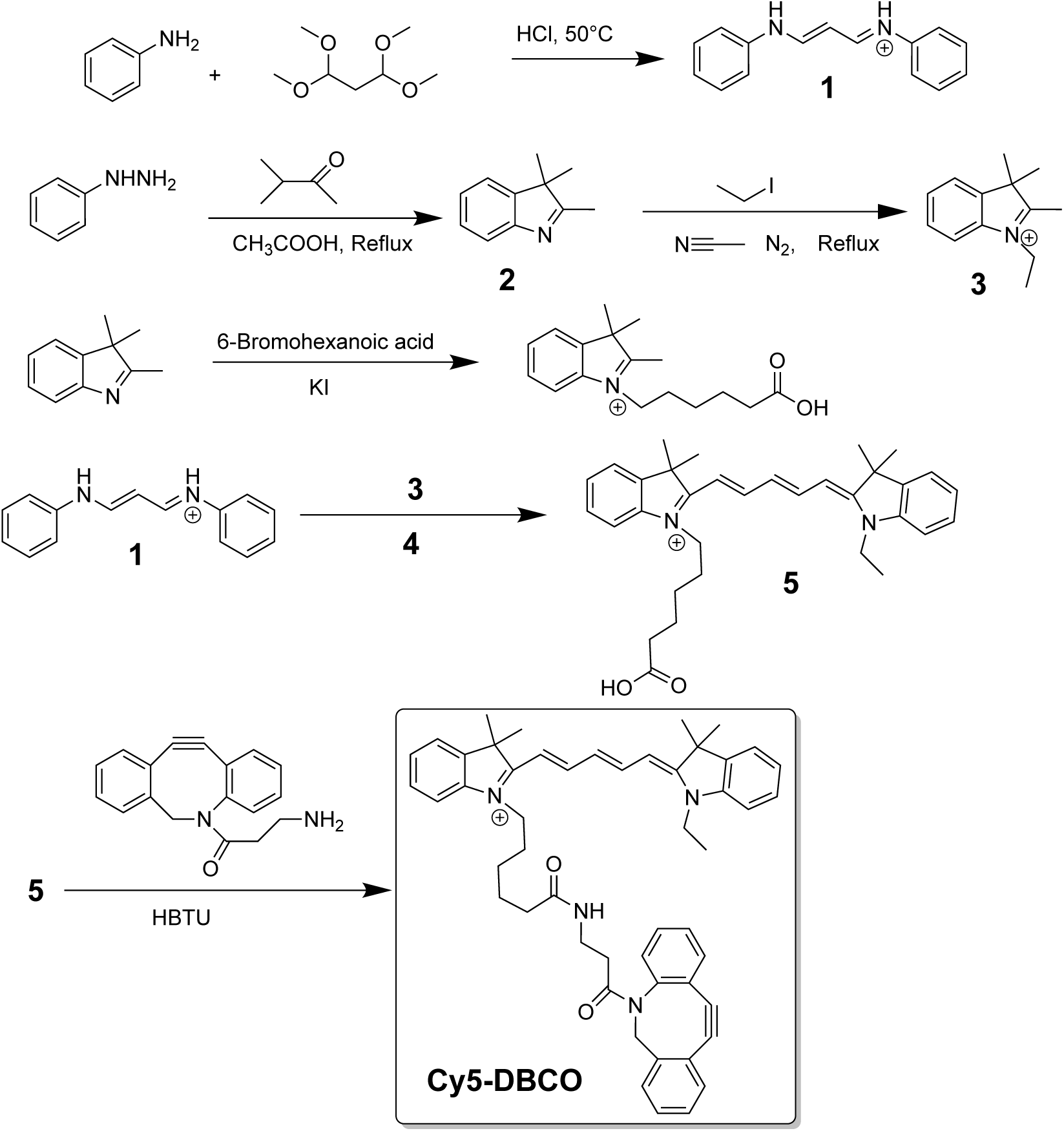
Synthetic scheme of preparation of Cy5-DBCO

### 2.3. Surface Engineering of BR-EVs for Mitochondrial Targeting

#### 2.3.1. Synthesis of the Mitochondrial-Targeting Cy5-DBCO Ligand

The Cy5-DBCO construct was synthesized through a modular route designed to assemble the asymmetric cyanine chromophore and subsequently append a strained DBCO group to enable copper-free click chemistry. The sequence began with formation of aldehyde intermediate **1** via acid-catalyzed condensation of 1,1,3,3-tetramethoxypropane with aniline. Proton-mediated generation of an iminium ion was followed by intramolecular ring closure, rearrangement, and deprotonation to afford the aromatic product.

Intermediate **2** was prepared by condensing phenylhydrazine with 3-methylbutanone in glacial acetic acid. The in-situ formation of an acetylated phenylhydrazine promoted imine formation, which upon protonation rearranged to a stabilized enamine that cyclized to furnish the heterocyclic scaffold. Subsequent N-alkylation with ethyl iodide in acetonitrile yielded quaternary indolium salt **3** via a canonical SN2 displacement. To generate the carboxyindoleninium salt 4, 2,3,3-trimethyl-3H-indole reacted with a transient acyl iodide prepared from 6-bromohexanoic acid and potassium iodide. Nucleophilic addition, internal rearrangement, and deprotonation produced a stabilized indoleninium intermediate. Intermediates **1**, **3**, and **4** were then condensed with malonaldehyde dianil to construct the asymmetric cyanine framework (compound **5**). Final amide coupling of **5** with DBCO-amine yielded the Cy5-DBCO fluorophore, preserving the spectral properties of Cy5 while incorporating a strained alkyne for SPAAC. The resulting dye, a delocalized lipophilic cation, demonstrated efficient mitochondrial localization in cultured cells and in vivo systems (images not shown), establishing its suitability for subsequent covalent modification of extracellular vesicles and real-time mitochondrial tracking.

**Scheme 2.**
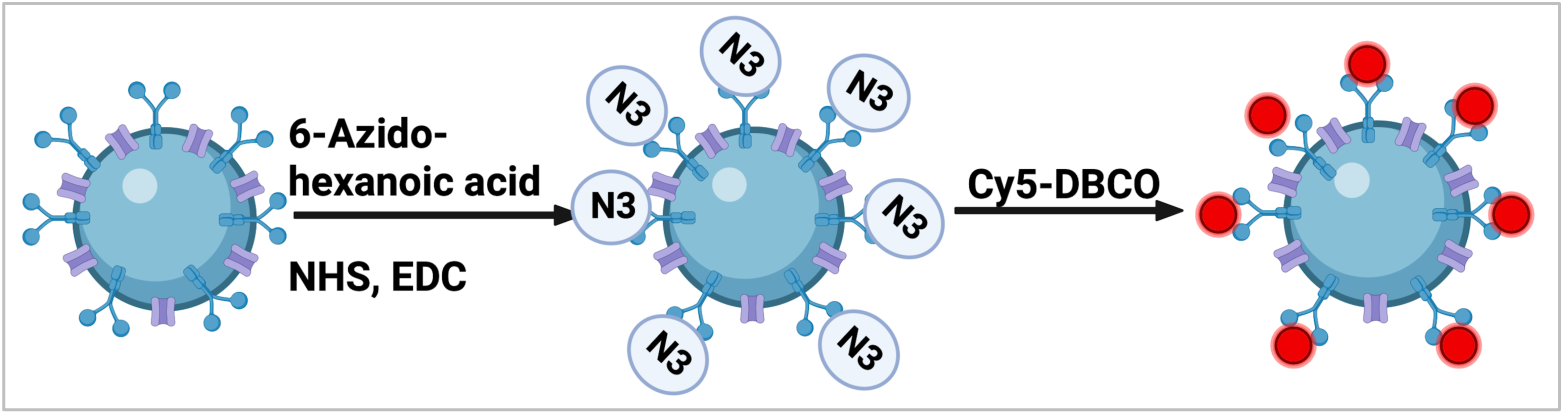
An illustration image of surface azide installation and copper-free conjugation to BR-EVs.

#### 2.3.2 Surface Azide Installation and Copper-Free Conjugation to BR-EVs

Given the pivotal role of exosomal surface architecture in dictating cellular uptake, subcellular routing, and cargo transfer fidelity, we developed a bioorthogonal functionalization strategy that preserves native vesicle morphology, membrane protein topology, and proteomic integrity. Primary amines on exosomal surface proteins—predominantly lysine residues on tetraspanins such as CD63 and CD81, as well as annexins (Anxa1–6, A8) identified in our proteomic dataset—served as selective anchor points for azide installation.

BR-EVs isolated from SD- and DSS-rats were reacted with activated 6-azido-hexanoic acid, generated by EDC/NHS coupling in MES buffer (Scheme 2). Excess carbodiimide was quenched with 2-mercaptoethanol, and the NHS ester was allowed to react with EVs surface amines in PBS (pH 7.2) at −4 °C for 12 h. Residual active esters were terminated with hydroxylamine, and azide-modified vesicles were purified by size-exclusion chromatography. Typical recoveries were 62–65% based on protein quantification.

Azido-functionalized BR-EVs were then incubated with Cy5-DBCO (250 µM) under light-protected conditions at −4 °C for 12 h. Following SPAAC conjugation, unbound fluorophore was removed by SEC, yielding comparable protein recovery. Fluorescence measurements demonstrated ∼0.1–0.2 nmol Cy5 per mg protein, corresponding to approximately 10–20% surface amine labeling. This density reflects steric constraints on the crowded exosomal membrane and was intentionally maintained at a sparse regime to avoid aggregation, perturbation of protein-mediated uptake pathways, and fluorophore self-quenching.

Post-modification quality control confirmed preserved particle integrity. Dynamic light scattering indicated unchanged hydrodynamic diameter (120–200 nm; PDI 0.15–0.25; p > 0.05), and TEM showed conserved cup-shaped morphology without aggregation. Untreated vesicles served as negative controls. These data confirm that SPAAC affords selective, copper-free conjugation that maintains vesicle biophysical properties and avoids oxidative damage associated with Cu-catalyzed click chemistry.

### 2.4. Complementary Time-Course Experiment: Uptake Kinetics and Mitochondrial Engagement of SPAAC-Engineered Exosomes

A key innovation of this work is the deployment of SPAAC to functionalize BR-EVs with a MTL (e.g. Cy5-DBCO). SPAAC offers a copper-free, bioorthogonal conjugation strategy that proceeds under mild aqueous conditions, avoids metal-mediated cytotoxicity, and preserves lipid membrane integrity. In contrast to conventional dye-loading approaches that rely on nonspecific lipid intercalation and exhibit labeling instability, SPAAC affords stable, covalent surface modification while maintaining vesicular architecture and bioactivity. This feature is essential for examining mitochondrial tropism in physiologically relevant settings, as the conjugate must withstand endosomal sorting, intracellular acidification, and organellar trafficking without perturbing natural signaling properties.

To rigorously interrogate uptake kinetics and subcellular fate, we conducted a longitudinal imaging study in primary cortical neurons obtained from wild-type rodents. Two SPAAC-engineered EV populations were evaluated in parallel: Cy5-DBCO-SD-EVs, isolated from normotensive rats and enriched in cytoprotective cargos, and Cy5-DBCO-DSS-EVs, derived from hypertensive donors bearing pro-oxidative and pro-inflammatory signatures. Neurons were treated with 2 μg mL⁻¹ of each construct, approximating physiologic EV abundances in neuro-inflammatory states. Live-cell confocal microscopy was performed at 6, 12, 24, and 48 hours post-treatment across biological triplicates, with ≥15 fields per coverslip, enabling quantitative assessment of exosomal uptake, Cy5 accumulation, mitochondrial localization, and mitochondrial morphology under nondisruptive imaging parameters.

Confocal z-stacks demonstrated progressive accrual of Cy5-labeled vesicles within neuronal somata and neurites, with clear punctate colocalization at the mitochondrial network. Yellow overlap between Cy5 and MitoTracker signals confirmed mitochondrial enrichment throughout the observation period, and no cytoplasmic dye diffusion was observed. Quantitative colocalization analysis yielded a Pearson’s coefficient of 0.82 ± 0.04, indicating strong linear association between EV and mitochondrial signals. Manders’ M1 values averaged 0.76 ± 0.06, consistent with high mitochondrial targeting efficiency, while M2 remained <0.1, demonstrating minimal nonspecific occupancy of mitochondrial space. Importantly, concurrent lysosomal staining (images not shown) revealed <5 % EVs–lysosome overlap at 48 h, indicating that SPAAC labeling does not redirect EVs toward degradative pathways.

To evaluate uptake dynamics independent of mitochondrial enrichment, Cy5 fluorescence intensities were quantified from neuronal cells following background correction and normalization to 6-hour values. Both EVs types displayed near-linear temporal accumulation: ∼1.4-fold at 12 h, ∼2.0-fold at 24 h, and ∼2.5-fold at 48 h (ANOVA, p < 0.001 for time; no significant interaction between EVs subtype and time, p = 0.78) (Table 1). The closely overlapping uptake profiles demonstrate that SPAAC modification neither impedes nor accelerates internalization and preserves canonical neuronal EVs uptake behavior. Notably, the absence of plateauing suggests sustained macropinocytic or phagocytic flux rather than saturable receptor-limited entry—an important property for therapeutic scenarios requiring extended mitochondrial cargo delivery.

**Table 1:**
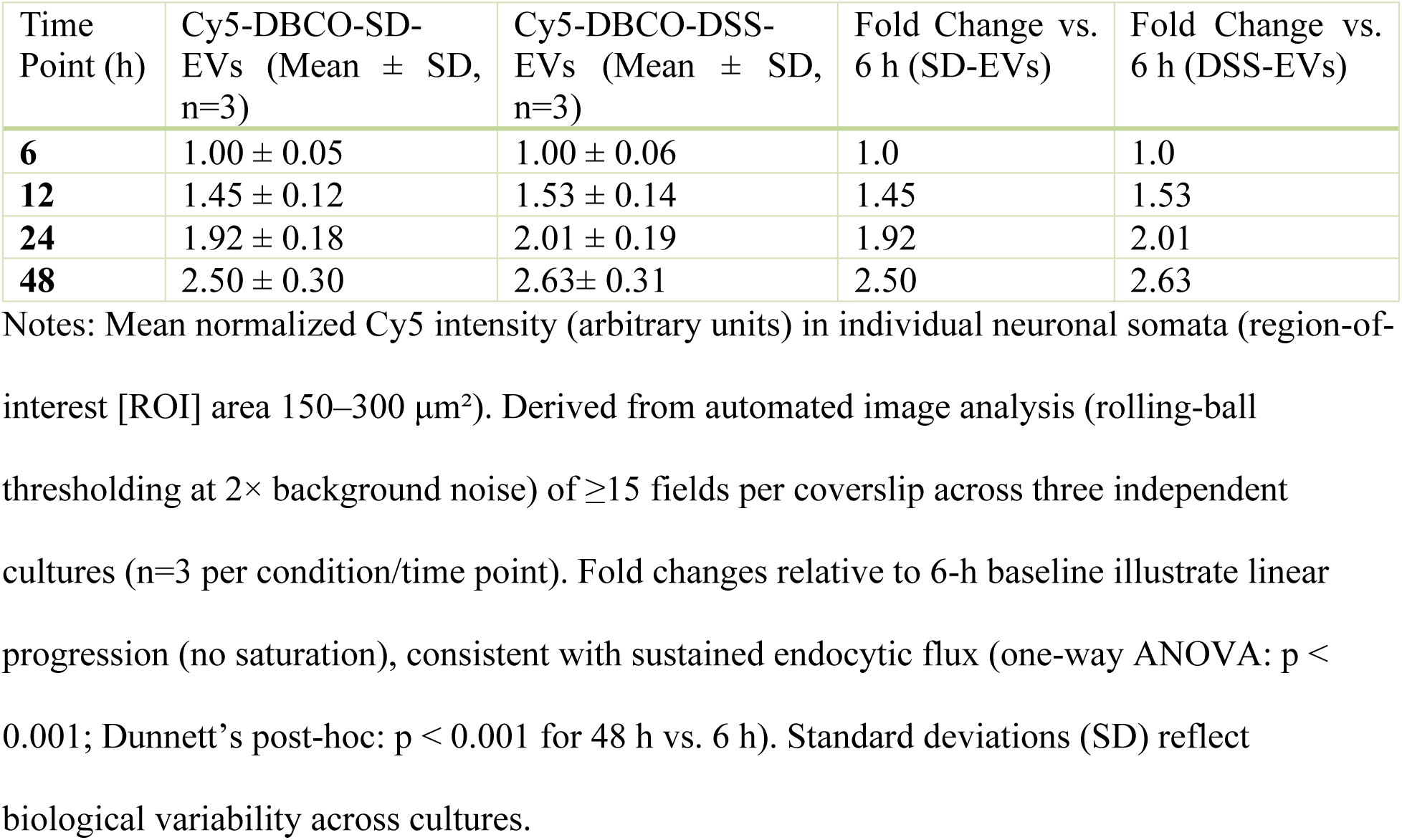
Time-Dependent Normalized Cy5 Fluorescence Intensity in Neuronal Somata.

#### 2.4.1. Differential Effects on Mitochondrial Morphology: SPAAC Labeling Preserves Organelle Architecture

To determine whether SPAAC-engineered EVs alter mitochondrial structure, we analyzed aspect ratio (AR) and form factor (FF)—quantitative parameters reflecting mitochondrial elongation and branching. Untreated neurons exhibited the characteristic tubular mitochondrial morphology (AR 4.5 ± 0.2; FF 0.40 ± 0.03), indicative of balanced fusion–fission dynamics. Exposure to Cy5-DBCO–SD-EVs maintained this architecture throughout the 48-hour observation period, with AR values remaining between 4.0 and 4.3 and FF between 0.40 and 0.46 (repeated-measures ANOVA, p > 0.3). These data confirm that SPAAC functionalization does not perturb mitochondrial ultrastructure or compromise dynamic equilibrium.

**Table 2.** Aspect Ratio (AR) Comparison.

| Time Point (h) | Cy5-DBCO-SD-EVs AR (mean ± SD) | Cy5-DBCO-DSS-EVs AR (mean ± SD) | Unlabeled DSS-EVs AR (mean ± SD) |
| --- | --- | --- | --- |
| 0 (baseline) | 4.5 ± 0.2 | 4.5 ± 0.2 | 4.5 ± 0.2 |
| 6 | 4.1 ± 0.3 | 4.0 ± 0.3 | 4.5 ± 0.3 |
| 12 | 4.0 ± 0.2 | 3.9 ± 0.2 | 3.9 ± 0.2 |
| 24 | 4.2 ± 0.3 | 3.0 ± 0.3 | 3.0 ± 0.3 |
| 48 | 4.3 ± 0.2 | 2.1 ± 0.2 | 2.1 ± 0.2 |

**Table 3.** Form Factor (FF) Comparison.

| **Time Point (h)** | **Cy5-DBCO-SD-EVs FF (mean ± SD)** | **Cy5-DBCO-DSS-EVs FF (mean ± SD)** | **Unlabeled DSS-EVs FF (mean ± SD)** |
| --- | --- | --- | --- |
| **0 (baseline)** | 0.40 ± 0.03 | 0.40 ± 0.03 | 0.40 ± 0.03 |
| **6** | 0.41 ± 0.03 | 0.40 ± 0.03 | 0.40 ± 0.03 |
| **12** | 0.46 ± 0.04 | 0.55 ± 0.02 | 0.55 ± 0.02 |
| **24** | 0.45 ± 0.03 | 0.72 ± 0.04 | 0.72 ± 0.04 |
| **48** | 0.44 ± 0.04 | 0.83 ± 0.05 | 0.83 ± 0.05 |

In contrast, neurons treated with Cy5-DBCO–DSS-EVs displayed progressive mitochondrial fragmentation, evidenced by a marked reduction in AR from 4.0 ± 0.2 to 2.1 ± 0.2 and a concomitant rise in FF from 0.40 ± 0.03 to 0.83 ± 0.05 by 48 hours. The shift toward shorter, rounded mitochondria signifies enhanced fission and structural collapse, hallmarks of oxidative stress and metabolic impairment. Notably, unlabeled DSS-EVs produced indistinguishable outcomes (AR/FF comparison, p > 0.9), demonstrating that the observed injury arises from EVs-borne pathogenic cargo rather than from the SPAAC modification itself. These findings establish that copper-free SPAAC labeling preserves vesicular bioactivity and allows faithful visualization of EVs-induced mitochondrial remodeling.

#### 2.4.2. Hypertensive EVs Cargo Drives Oxidative Stress and Mitochondrial Fragmentation Independent of SPAAC Labeling

To dissect the mechanistic basis of DSS-EV-induced mitochondrial fragmentation observed in the AR/FF analysis, we integrated SPAAC-enabled colocalization phenotyping with prior functional data from unlabeled EVs. These complementary assays in unlabeled SD- and DSS-EVs revealed a time-dependent mtROS pattern: both induced initial accrual, but SD-EV-treated neurons showed a sharp decline by 48 h, whereas DSS-EV exposure sustained a 2-fold elevation in mtROS intensity (MitoProbe), indicative of prolonged pro-oxidative signaling. This was coupled to robust inflammatory activation in DSS-EV cultures, including 2.3-fold TNFα, 3.7-fold IL1β, and 1.4-fold NF-κB mRNA upregulation (qPCR; p < 0.01 vs. SD-EVs), alongside 1.3-fold c-Fos elevation and chemokine surges (CCL2 2.4-fold, CCL5 2.1-fold, CCL12 4.2-fold; p < 0.001), evoking NOX2-biased NADPH oxidase priming and immune mobilization. ^37^

These oxidative and inflammatory phenotypes are strongly supported by our proteomic analysis of native EVs (Table S1). DSS-EVs were enriched in oxidative stress–associated chaperones such as Hspa2 and Hspa1l, and Nrf2-linked detoxification enzymes including Gstm7 and Gsta3, highlighting vesicular mobilization of maladaptive antioxidant and protein-refolding machinery under hypertensive stress. DSS-EVs also contained mitochondrial respiratory chain proteins (Ndufs2, Ndufa5) in altered abundance, suggesting metabolic repurposing toward pro-oxidant signaling. Mitochondrial signatures—unique Acsbg1 (β-oxidation ligase) and Rab GTPases (Rab1b/8b/10/13) for fission-biased trafficking—further evoked mitophagic debris packaging, absent in SD-EVs (13 unique proteins, primarily cytoskeletal). In parallel, inflammatory cargo including DSS-unique complement components C3 and C4, APP, and shared annexins (Anxa2/5/6) indicates a neuroimmune-amplifying phenotype, with annexins fueling eicosanoid release via phospholipase A₂ modulation. Shared phospho-ATPases (Atp1a1/2/3 at T366/T674, T364, T233/T356) and vesicle transport protein Bnip1 imply dysregulated Ca²⁺ homeostasis and ER-mitochondria-endosome crosstalk, priming fragmentation. The detection of these Rab GTPases and mitochondrial fragments in DSS-EVs implicates stress-induced mitovesicle release, consistent with enhanced turnover during hypertension.

These findings reveal a coordinated oxidative–inflammatory–mitochondrial stress program selectively exported by hypertensive EVs, sufficient to disrupt mitochondrial integrity in recipient neurons. SPAAC-engineered DSS-EVs faithfully recapitulate these effects, confirming that bioorthogonal click modification preserves native vesicle function and molecular identity. This fidelity enables high-resolution tracking of subcellular vesicle trafficking and provides a powerful platform for dissecting EV-mediated redox-immune signaling and mitochondrial vulnerability in hypertensive neuroinflammation.

#### 2.4.3. Uptake Mechanisms: Bulk Endocytosis as the Predominant Entry Route; SPAAC Modification Preserves Internalization Dynamics

Live-cell confocal imaging over 48 hours revealed a progressive, time-dependent accumulation of Cy5 fluorescence within neuronal cells, indicating sustained internalization of SPAAC-engineered BR-EVs (Figure 1–2). Quantitative analysis showed a 2.5-fold increase in intracellular Cy5 intensity by 48 h relative to the 6 h baseline, with no evidence of saturation (Table 1). The continuous increase suggests that uptake occurs predominantly through high-capacity, dynamin-dependent bulk endocytosis—such as macropinocytosis or phagocytic-like engulfment—rather than through saturable, receptor-mediated mechanisms. The uptake kinetics were consistent across three independent neuronal cultures (≥15 imaging fields each), underscoring reproducibility.

**Figure 1.**
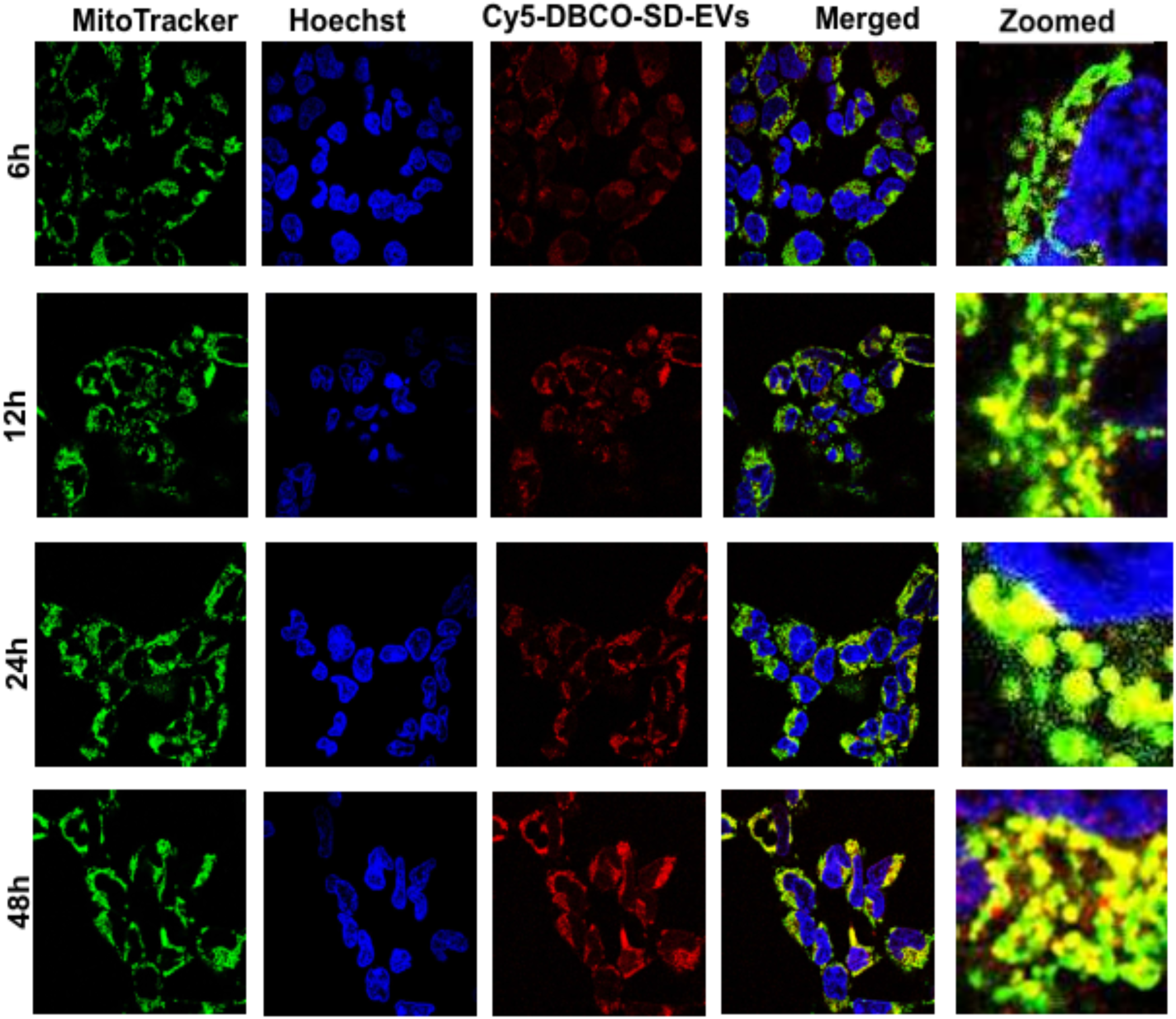
Representative confocal fluorescent images show primary neuron cells incubated with Cy5-DBCO-SD-EVs (2 µg, red), co-stained with MitoTracker (80 nM, green) and Hoechst 33242 (0.1 µL/mL, blue), with merged images appearing yellow. The time-course study was conducted at 6, 12, 24 and 48 hours. Images were captured using a confocal laser scanning fluorescent microscope with a 60x objective lens in non-FBS, non-phenol red media.

**Figure 2.**
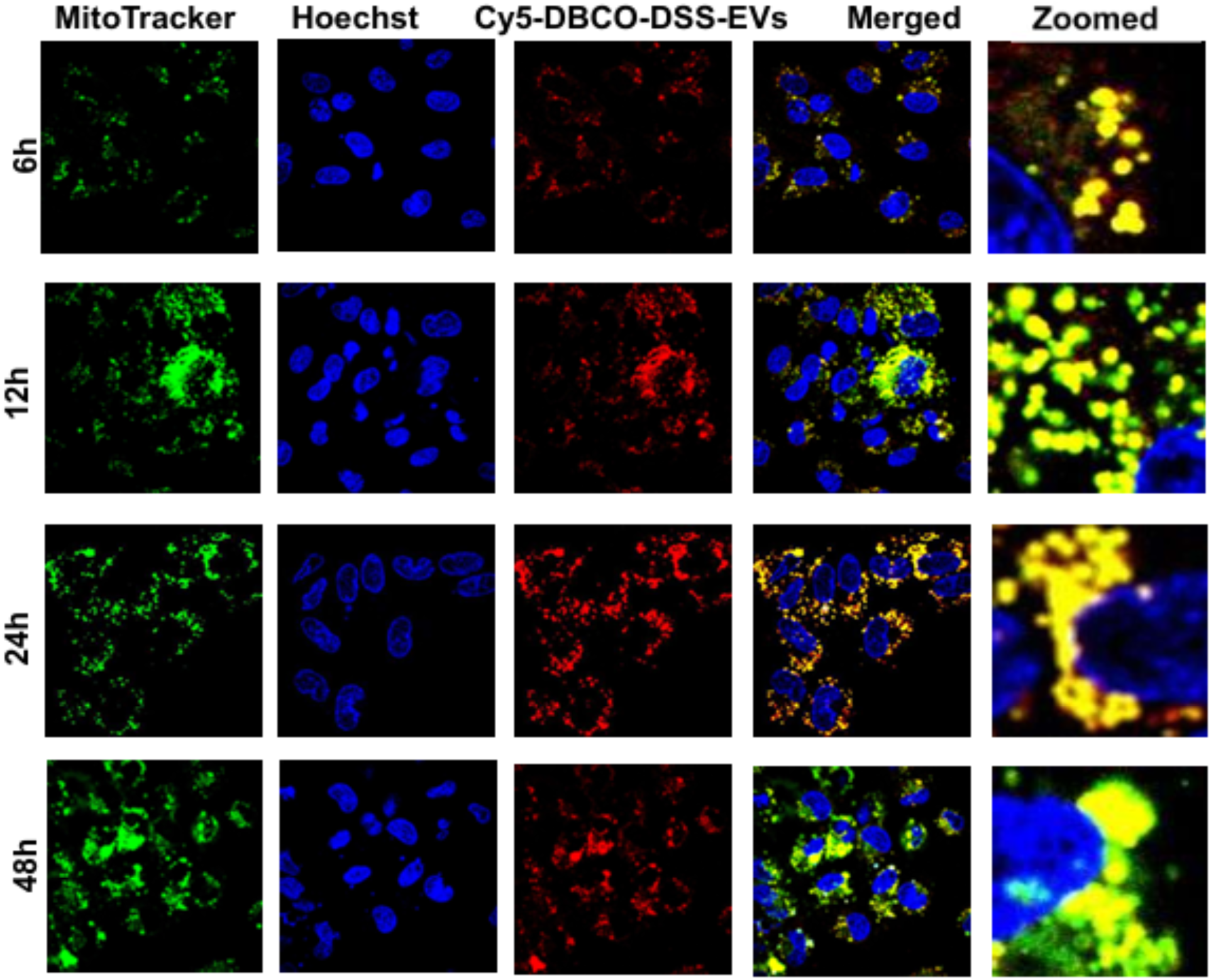
Representative confocal fluorescent images show primary neuron cells incubated with Cy5-DBCO-DSS-EVs (2 µg, red), co-stained with MitoTracker (80 nM, green) and Hoechst 33242 (0.1 µL/mL, blue), with merged images appearing yellow. The time-course study was conducted at 6, 12, 24 and 48 hours. Images were captured using a confocal laser scanning fluorescent microscope with a 60× objective lens in non-FBS, non-phenol red media.

Proteomic profiling of unlabeled SD- and DSS-derived EVs (760 proteins total; Table S1-S3) supported this interpretation. Both subtypes contained a conserved suite of endocytic and trafficking proteins, including Dynamin-1 (Dnm1), clathrin heavy and light chains (Cltc, Clta), VAMPs (1/2/3), NSF, and multiple Rab GTPases (Rab1A/5A/7A/11A), consistent with vesicle scission and intracellular trafficking through canonical dynamin-dependent pathways. Annexins (Anxa1–6, A8), known for inducing membrane curvature, further corroborated the observed non-saturable kinetics. DSS-specific Rab isoforms (Rab1B, Rab8B, Rab10, Rab13) may influence post-endocytic trafficking toward mitochondria but do not alter entry dynamics, as overall uptake rates remained statistically equivalent between SD- and DSS-EVs (p = 0.78).

Z-stack confocal imaging confirmed progressive intracellular accumulation of Cy5-DBCO–labeled EVs with increasing colocalization to the mitochondrial network over time. Hoechst nuclear counterstaining demonstrated intact nuclear morphology, excluding cytotoxic effects. Statistical analysis validated the time-dependent uptake (one-way ANOVA, p < 0.001; Dunnett’s post hoc, p < 0.001 for 48 h vs. 6 h).

The SPAAC conjugation process, involving < 20% surface amine occupancy, was specifically designed to preserve native endocytic ligands and vesicular membrane functionality. The observed sustained uptake, together with unaltered neuronal morphology, indicates that copper-free SPAAC chemistry does not perturb essential recognition motifs or internalization pathways.

Pharmacological inhibition assays at 24 h further elucidated the entry mechanisms. Treatment with dynasore (80 μM), a dynamin GTPase inhibitor, reduced Cy5-DBCO-BR-EVs uptake by 67 ± 8% for SD-EVs and 69 ± 10% for DSS-EVs (one-way ANOVA, p < 0.001; Dunnett’s post hoc, p < 0.01 vs. vehicle), confirming dynamin-dependent vesicle scission as a dominant pathway. In contrast, nystatin (50 μM), which disrupts caveolae-mediated endocytosis, produced negligible inhibition (1 ± 4% for SD-EVs; 3 ± 6% for DSS-EVs; p > 0.05), despite the presence of Cavin1 in the shared proteome, suggesting that caveolar routes play only a minor role. Two-way ANOVA detected no significant interaction between EVs subtype and inhibitor treatment (p = 0.78), confirming that SD- and DSS-derived EVs share a common internalization mechanism. Cell viability remained ≥80% across all groups, excluding cytotoxic confounds (Table 4).

**Table 4.** Inhibitor Effects on Cy5-DBCO-BR-EVs Uptake in Primary Neurons at 24 h.

| Inhibitor | Concentration | EV Variant | % Uptake Reduction (Mean ± SD, n=3) | p-value (vs. Vehicle) | Cell Viability (% MTT Assay, Mean ± SD) |
| --- | --- | --- | --- | --- | --- |
| Dynasore | 80 μM | Cy5-DBCO-SD-EVs | 67 ± 8% | <0.01 | 83 ± 4% |
| Dynasore | 80 μM | Cy5-DBCO-DSS-EVs | 69 ± 10% | <0.01 | 81 ± 5% |
| Nystatin | 50 μM | Cy5-DBCO-SD-EVs | 1 ± 4% | >0.05 | 83 ± 3% |
| Nystatin | 50 μM | Cy5-DBCO-DSS-EVs | 3 ± 6% | >0.05 | 84 ± 4% |
| DMSO (vehicle) | 0.1% | Both EVs | 0 ± 3% | N/A | 85 ± 2% |

These findings establish bulk endocytosis as the predominant neuronal entry route for BR-EVs and demonstrate that SPAAC modification preserves intrinsic internalization dynamics. The copper-free reaction maintains functional integrity of surface ligands critical for neuronal recognition and uptake, reinforcing the biocompatibility and mechanistic fidelity of this chemical engineering strategy.

#### 2.4.4. FRAP Analysis Reveals Altered Vesicle Mobility upon Mitochondrial Docking

To characterize the dynamic behavior of Cy5-DBCO-BR-EVs following their association with neuronal mitochondria, the FRAP assay was conducted on live primary neurons 24 hours after incubation with Cy5-DBCO–labeled BR-EVs (Table 5). FRAP provides a quantitative measure of molecular mobility by tracking fluorescence restoration within a bleached region of interest (ROI); slower or incomplete recovery reflects constrained diffusion and stable anchoring, whereas rapid recovery indicates enhanced mobility and weaker association.

**Table 5:**
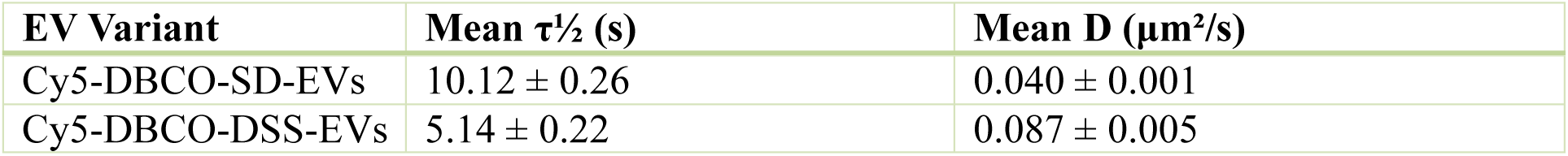
FRAP-Derived Half-Recovery Times (τ½) and Diffusion Coefficients (D) for Cy5-DBCO-BR-EVs.

| EV Variant | Mean $\tau_{1/2}$ (s) | Mean D ($\mu\text{m}^2/\text{s}$) |
| --- | --- | --- |
| Cy5-DBCO-SD-EVs | $10.12 \pm 0.26$ | $0.040 \pm 0.001$ |
| Cy5-DBCO-DSS-EVs | $5.14 \pm 0.22$ | $0.087 \pm 0.005$ |

Circular ROIs (2–3 μm in diameter) were defined at sites where Cy5-labeled EVs puncta colocalized with MitoTracker Green–labeled mitochondria. Photobleaching was performed using a 633 nm laser at full power for five iterations (1.27 μs per pixel dwell time), followed by fluorescence recovery imaging for five minutes at 5% laser power (one frame every three seconds). Half-recovery time (τ½) was determined as the interval required to regain 50% of the pre-bleach fluorescence, and diffusion coefficients (D) were computed using the relation D = w²/(4τ½), where w is the ROI radius (2.5 μm midpoint). Image J’s FRAP analysis plug-in was employed for curve fitting, and a minimum of 20 ROIs per condition were analyzed from three independent neuronal cultures. Statistical significance was assessed using unpaired t-tests (p < 0.05).

Normotensive Cy5-DBCO-SD-EVs exhibited a mean τ½ of 10.12 ± 0.26 s and a corresponding diffusion coefficient of 0.040 ± 0.001 μm²/s, consistent with restricted lateral mobility and stable mitochondrial docking. In contrast, hypertensive Cy5-DBCO–DSS-EVs demonstrated a markedly shorter τ½ of 5.14 ± 0.22 s and an increased diffusion coefficient of 0.087 ± 0.005 μm²/s—representing an approximate 50% reduction in τ½ and a two-fold elevation in D relative to SD-EVs. Back-calculated values confirmed internal consistency (e.g., D ≈ (2.5)² / [4 × 10.12] = 0.039 μm²/s for SD-EVs).

The diffusion coefficients observed (0.01–0.1 μm²/s) align with reported values for vesicles engaged in near-organelle interactions and are notably slower than those of freely diffusing cytosolic proteins (>1 μm²/s). These findings suggest that both EV populations establish physical associations with neuronal mitochondria; however, BR-EVs derived from hypertensive donors display enhanced motility and reduced docking stability. Such behavior implies that hypertension-associated cargo may weaken EVs–mitochondria tethering, potentially reflecting altered surface protein composition or diminished affinity for mitochondrial docking machinery.

### 2.5. Neuroinflammatory Effects and Glial Activation Analysis

A central objective of this study was to rigorously evaluate whether the copper-free SPAAC chemistry—specifically Cy5-DBCO conjugation—elicits neuroinflammatory responses when applied to BR-EVs. As glial activation represents an early and sensitive indicator of neuroimmune perturbation, we focused on canonical astrocytic and microglial markers: glial fibrillary acidic protein (GFAP), ionized calcium-binding adapter molecule 1 (IBA1), and cluster of differentiation 68 (CD68). Upregulation of these markers typifies reactive gliosis, a process associated with oxidative stress, cytokine secretion (e.g., IL-1β, TNF-α), and compromised blood–brain barrier (BBB) integrity. Thus, demonstrating the absence of glial activation following SPAAC modification provides a stringent benchmark of biocompatibility and translational safety for exosome-based therapeutics.

Quantitative analyses targeted neuroinflammation-prone regions, including the dentate gyrus (DG), CA1, and CA3 subfields of the hippocampus, and cortical layers II–V. Fluorescence intensity was quantified in ImageJ (v1.53) across anatomically defined ROIs, normalized to area and background. Two-way ANOVA (treatment × region) with Tukey’s post hoc testing (α = 0.05, GraphPad Prism v9) determined statistical significance.

Cy5-DBCO–functionalized SD-EVs produced no measurable neuroimmune activation in vivo. Quantitative immunohistochemistry demonstrated that GFAP, IBA1, and CD68 signals in Cy5-DBCO–EV–treated animals were indistinguishable from PBS controls across all brain regions examined. In the dentate gyrus, GFAP immunoreactivity exhibited a fold change of 1.05 ± 0.12 (F(1,20) = 0.42, p = 0.52), IBA1 of 1.08 ± 0.10 (F(1,20) = 0.65, p = 0.43), and CD68 of 1.02 ± 0.09 (F(1,20) = 0.18, p = 0.68), with similarly non-significant results in CA1, CA3, and cortex (all p > 0.05). All values remained within 5–10% of baseline, confirming that SPAAC-engineered EVs do not provoke astroglial or microglial activation.

This immunologically silent profile stands in sharp contrast to the pronounced GFAP and IBA1 upregulation reported for non-bioorthogonal nanomaterials, including uncoated quantum dots, underscoring the exceptional biocompatibility and inertness of SPAAC chemistry. The absence of gliosis further indicates that SPAAC modification preserves both the immunological quiescence and structural integrity of BR-EVs. Performed entirely under aqueous, neutral-pH, copper-free conditions, SPAAC avoids the redox stress, membrane perturbation, and cytotoxic byproducts characteristic of CuAAC and other reactive coupling chemistries. These findings establish Cy5-DBCO labeling as a robust and fully bioorthogonal strategy for tracking mitochondrial-targeted EVs in vivo without eliciting adverse neuroimmune responses.

**Figure 3.**
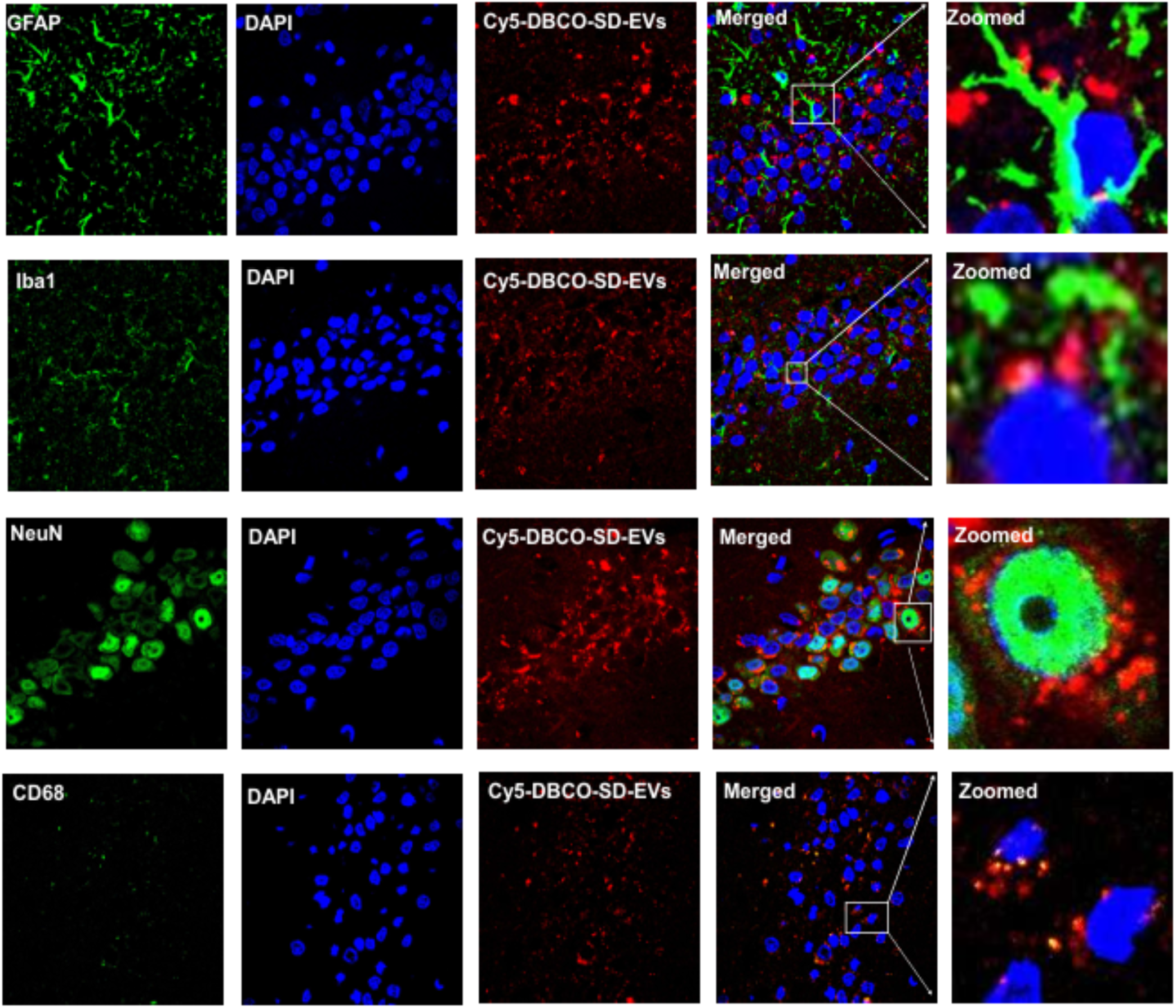
Evaluation of neuroinflammatory response following intracerebroventricular (ICV) administration of Cy5-DBCO-SD-EVs. Adult SD rats received Cy5-DBCO-SD-EVs or vehicle (PBS), and brain sections were immunostained for GFAP, IBA1, CD68, and NeuN to assess astrocytic, microglial, and neuronal markers in the hippocampus and cortex. Confocal images were acquired to quantify fluorescence intensity of glial markers, normalized to DAPI-positive nuclei, and compared between Cy5-DBCO-SD-EV–treated and PBS control groups (control images not shown).

This in vivo validation was intentionally confined to Cy5-DBCO-SD-EVs, derived from normotensive donors, to isolate the biocompatibility of the SPAAC process itself. *In vivo* testing of Cy5-DBCO-DSS-EVs was not performed, as proteomic analysis of unlabeled DSS-EVs revealed extensive enrichment in oxidative and inflammatory mediators—including glutathione S-transferases (GSTs), complement components (C3, C4), and Rab-family GTPases—signifying an intrinsic pro-inflammatory and redox-active cargo profile. Subsequent *in vitro* experiments confirmed that DSS-EVs induced mitochondrial fragmentation, elevated mtROS (MitoProbe), and astroglial activation in neuron–astrocyte co-cultures—effects absent in SD-EVs. Introducing such pathologic cargo into the *in vivo* system would have confounded interpretation of SPAAC’s safety and masked its true biocompatibility. Thus, the decision to exclude Cy5-DBCO-DSS-EVs from animal studies was deliberate, ensuring that the observed outcomes directly reflected the chemical inertness of the SPAAC reaction rather than the biological toxicity of hypertensive EVs cargo.

These findings establish that SPAAC-engineered exosomes achieve precise functional labeling with exceptional biocompatibility. Cy5-DBCO conjugation yields ∼10–20% labeling efficiency while maintaining hydrodynamic diameters (120–200 nm by DLS) and canonical cup-shaped morphology (30–150 nm by TEM). The resulting exosomes retain structural and immunological stability, enabling real-time tracking of mitochondrial interactions in vivo without initiating glial activation. In the broader context, this finding addresses a key translational bottleneck: the challenge of functionalizing exosomes for targeted brain delivery while avoiding neuroinflammatory side effects—a property that distinguishes SPAAC-engineered BR-EVs as a clinically viable platform for precision neurotherapeutics.

### 2.6. Biodistribution and Clearance of Cy5-DBCO–Labeled SD-EVs

Following ICV administration, Cy5-DBCO-SD-EVs exhibited a predominantly central nervous system (CNS)-restricted distribution, consistent with cerebrospinal fluid (CSF)-driven dissemination throughout periventricular, hippocampal, and cortical regions. This pattern reflects the strong biocompatibility of the SPAAC modification, as copper-free click conjugation preserves vesicular structure and surface chemistry under mild, aqueous conditions, avoiding the oxidative or membrane-disruptive side effects often associated with metal-catalyzed reactions. The low Cy5 labeling density (∼0.1–0.2 nmol mg⁻¹ protein) was sufficient for stable fluorescence tracking while maintaining native size and colloidal stability.

A small fraction of Cy5 fluorescence was detected in peripheral organs, most prominently in the liver, with faint or negligible signals observed in the lungs, kidneys, intestine, adrenal gland, and heart. The modest hepatic signal likely reflects physiological clearance via glymphatic and meningeal lymphatic drainage into systemic circulation, followed by uptake by hepatic sinusoidal cells and Kupffer macrophages—the primary clearance route for circulating vesicles. This pattern parallels the transient systemic appearance of endogenous exosomes and suggests normal physiological turnover rather than nonspecific leakage or inflammatory accumulation.

**Figure 4:**
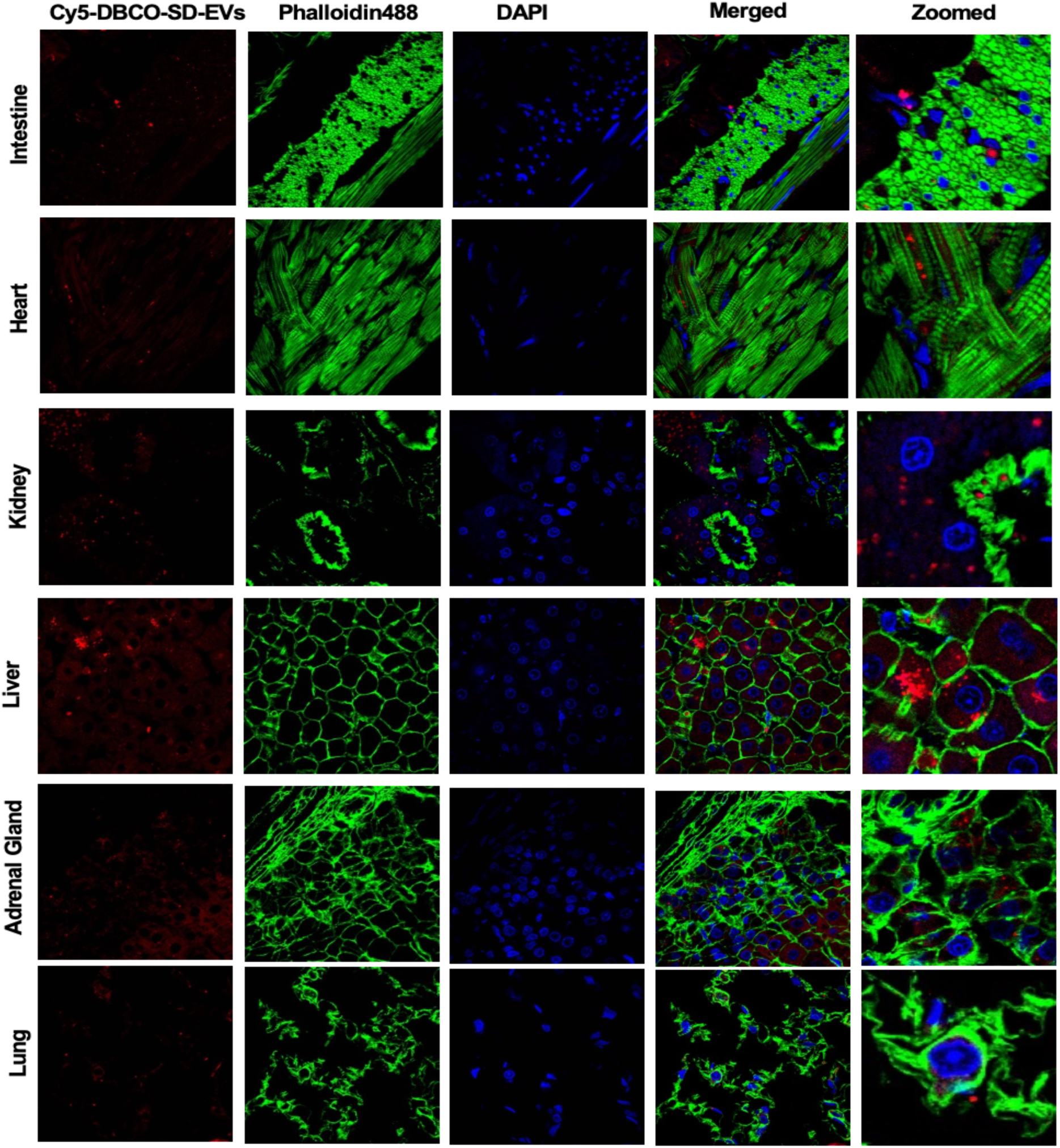
Representative confocal images showing the biodistribution of Cy5-DBCO–labeled SD-EVs. Confocal fluorescence images of intestine, liver, lung, kidney, adrenal gland, and heart tissues collected 24 h after ICV administration of Cy5-DBCO–SD-EVs (5 μg total protein per rat). Cy5 fluorescence (red) indicates exosome localization, DAPI (blue) stains cell nuclei, and phalloidin (green) labels the actin cytoskeleton. Images were acquired using a 60× objective on a confocal microscope.

Mechanistically, CSF-borne EVs enter perivascular glymphatic flow driven by arterial pulsations and aquaporin-4–mediated water flux at astrocytic endfeet, facilitating gradual redistribution through cervical lymphatics and venous return. Semi-quantitative fluorescence analysis indicated that approximately two-thirds of total Cy5 signal remained within the brain 24 h post-injection, confirming robust CNS retention and validating ICV administration as an efficient method for bypassing the blood–brain barrier. The absence of histological abnormalities or glial activation further supports that SPAAC modification does not alter vesicle biodistribution or compromise tissue integrity.

These results demonstrate that SPAAC-engineered SD-EVs preserve the natural trafficking and clearance characteristics of native exosomes. The combination of high CNS retention limited peripheral exposure, and lack of detectable toxicity underscores the biocompatibility and translational potential of this copper-free click chemistry approach for safe, traceable exosome-based brain delivery.

## 3. Materials and Methods

### 3.1. Animal Models and Comparative Analysis

Adult SD and DSS rats were obtained from Charles River Laboratories (Wilmington, MA, USA) and bred in our colony to generate offspring. Animals were housed under controlled conditions (20–24 °C, 12 h light/dark cycle) with ad libitum access to food and water. They were used for ICV injection, brain-derived EV isolation, and primary neuronal culture preparation, as described below. All procedures were approved by the IACUC at Michigan Technological University.

To induce hypertension, six-week-old male DSS rats were fed a high-salt diet (4% NaCl) for six consecutive weeks, resulting in chronic hypertension (systolic blood pressure >160 mmHg, confirmed by tail-cuff plethysmography). EVs from hypertensive DSS rats and age-matched normotensive SD rats maintained on standard chow (0.4% NaCl) were then isolated as described previously^39^. Comparative structural analyses of pooled BR-EVs (n = 6 rats per group) were performed using TEM and DLS, with statistical comparisons conducted using unpaired t-tests (α = 0.05).

### 3.2. Isolation of BR-EVs

Whole rat brains (excluding the cerebellum) were maintained in phosphate-buffered saline (PBS) and diced into small fragments on ice. Portions of tissue (200 mg each) were then distributed into wells of a 6-well plate, with each well containing 2 mL of Hibernate E medium (Gibco). The fragments were gently homogenized until uniformly sized, approximating 2 mm in each dimension (cube-like). To each well, 40 μL of Collagenase D (Sigma-Aldrich) and 4 μL of DNase I (Sigma-Aldrich) were introduced, yielding final concentrations of 2 mg/mL for Collagenase D and 40 U/mL for DNase I. The plate was promptly placed in a 37 °C incubator with gentle shaking (70 rpm) for 30 min. Post-incubation, the plate was returned to ice, and a cocktail of protease and phosphatase inhibitors (Sigma-Aldrich) was added to every well. The mixture, comprising tissue remnants and Hibernate E, was allowed to pass by gravity through a 70 μm sterile cell strainer into a 50 mL conical tube. Residual material in the wells was recovered by rinsing with an additional 1–2 mL of PBS.

The strained suspension underwent sequential differential centrifugation at 4 °C: first at 300 × g for 10 min, followed by 2,000 × g for 20 min, and then 10,000 × g for 30 min. Supernatants from these steps were retrieved and passed through a 0.22 μm filter (Grainger Industrial Supply) to eliminate residual debris. The clarified filtrates were carefully overlaid onto 4 mL cushions of 30% (w/v) sucrose in PBS and subjected to ultracentrifugation at 100,000 × g for 90 min at 4 °C using an Optima XL-90 ultracentrifuge equipped with an SW 28 swinging-bucket rotor (Beckman-Coulter). After discarding the upper supernatant, the sucrose interface (∼5 mL) and pelleted material were harvested. This fraction was further purified by a second round of ultracentrifugation under identical conditions (100,000 × g, 90 min, 4 °C), after which the resultant pellet was gently resuspended in 50 μL of ice-cold PBS for downstream applications.

For total protein assessment, BR-EVs were lysed in radioimmunoprecipitation assay (RIPA) buffer supplemented with 0.5% phenylmethylsulfonyl fluoride (PMSF; Sigma-Aldrich). Briefly, an equal volume of this lysis buffer was added to the vesicle suspension, followed by sonication (three cycles of 5 s each). The samples were then incubated on ice for 20 min, with intermittent pipetting every 5 min to ensure complete disruption. Protein concentrations were subsequently measured using the Bradford assay (Sigma-Aldrich).

### 3.3. DLS Characterization of Exosomes

Resuspended exosome preparations were adjusted with Dulbecco’s phosphate-buffered saline (DPBS) to a target protein level of 0.01 μg/μL. These adjusted suspensions were loaded into low-volume cuvettes, where particle size profiles were evaluated on a Malvern Zetasizer Nano series instrument. To minimize settling of larger aggregates, the capped cuvettes were inverted gently before measurement. Triplicate acquisitions were performed for every preparation.

### 3.4. TEM Evaluation of Exosome Ultrastructure

A 30 μL aliquot of the exosome resuspension was combined with 30 μL of 2% paraformaldehyde (PFA) and incubated for 5 minutes. From this mixture, 5 μL was deposited onto the carbon-coated face of a glow-discharged grid and permitted to settle for 1 minute. The liquid was then delicately wicked away using pristine filter paper. For contrast enhancement, 3 μL of freshly prepared 2% uranyl acetate (UA) was applied to the grid surface for 30 seconds, after which surplus stain was removed by blotting with fresh filter paper. Following air drying, the preparations were imaged on an FEI 200 kV Titan Themis scanning transmission electron microscope.

### 3.5. Surface Engineering and Functionalization of BR-EVs

Surface modification of BR-EVs exploited primary amines on exosomal proteins for covalent attachment of functional groups. Azido-functionalization was achieved by activating 6-azido-hexanoic acid (10 mM) with 1-ethyl-3-(3-dimethylaminopropyl)carbodiimide (EDC; 20 mM) and N-hydroxysuccinimide (NHS; 5 mM) in 2-(N-morpholino)ethanesulfonic acid (MES) buffer (0.1 M, pH 6.0) for 15 min at room temperature. Excess EDC was quenched with 10 mM 2-mercaptoethanol for 10 min, and the activated NHS ester (1 mM final) was added to BR-EVs (1 mg/mL protein) in PBS (pH 7.2) for 12 h at 4°C with gentle rotation. Unreacted esters were quenched with 50 mM hydroxylamine (pH 7.4) for 1 h at 4°C. Azido-BR-EVs were purified by size-exclusion chromatography (SEC) on Sepharose CL-4B columns (bed volume 10 mL, equilibrated with PBS), eluting in 1-mL fractions and collecting vesicle-enriched fractions (void volume) based on Bradford assay.

Subsequent Cy5 labeling employed SPAAC click chemistry. Azido-BR-EVs (0.5 mg/mL) were incubated with dibenzo-cyclooctyne (DBCO)-Cy5 conjugate (250 μM final) in PBS (pH 7.4) for 12 h at 4°C in the dark to prevent photobleaching. Unbound dye was removed by SEC as above, yielding 62–65% recovery (determined by protein quantification). Labeling efficiency was assessed by fluorescence spectrophotometry (excitation 650 nm, emission 670 nm; SpectraMax i3x reader), normalized to total protein via BCA assay, targeting ∼1 μmol surface amines per mg protein. Untreated BR-EVs served as negative controls. Post-modification integrity was verified by repeat DLS and TEM.

### 3.6. Cell Culture, Imaging, and Functional Assays

Primary neurons were isolated from brains of one-day-old SD pups and plated at 5×10^4 cells/cm² on poly-D-lysine-coated glass coverslips. Cells were incubated in neurobasal medium supplemented with B27 (2%) serum and 1% penicillin-streptomycin, and maintained at 37°C incubator with 5% CO₂ for 7∼12 days before using for experiments. For uptake studies, cells were incubated with Cy5-DBCO-labeled BR-EVs (2–10 μg/mL total protein) from SD or DSS rats. Active mitochondria were labeled by pre-incubating with 200 nM MitoTracker Green FM for 30 min at 37°C, followed by three PBS washes. Nuclei were counterstained with 0.1 μL/mL Hoechst 33242. Lysosomal targeting was assessed using 100 nM LysoTracker Green (optional co-stain).

Live-cell imaging was conducted on a confocal laser scanning microscope equipped with a 37°C/5% CO₂ environmental chamber and 60× oil-immersion objective (NA 1.4). Z-stack projections (0.5-μm slices, 1024×1024 pixels, 0.1-μm dwell time, 4× line averaging) were acquired from 20–30 random fields (≥50 neurons/field) per coverslip using sequential excitation (488 nm for MitoTracker/LysoTracker, 633 nm for Cy5, 405 nm for Hoechst) to minimize crosstalk. Time-course imaging occurred at 6, 12, 24, and 48 h post-incubation in non-fetal bovine serum (FBS), non-phenol red medium. Colocalization was quantified using Pearson’s correlation and Manders’ overlap coefficients via Fiji/ImageJ colocalization plugin (thresholded at 2× background).

Fluorescence intensity in neuronal cells (ROI: 150–300 μm², automated via DAPI-guided segmentation) was analyzed using rolling-ball background subtraction (radius 50 pixels) and normalized to 6-h baseline. Mitochondrial morphology was segmented semi-automatically in ImageJ (thresholded, watershed-separated) to compute aspect ratio (AR; major/minor axis) and form factor (FF; 4π × area / perimeter²) from ≥200 mitochondria per condition/time point across n=3 cultures. Oxidative stress was evaluated after 24-h exposure: total RNA was extracted (TRIzol), reverse-transcribed, and qRT-PCR performed for CYBA and CYBB (normalized to GAPDH; ΔΔCt method). Mitochondrial ROS was measured using 5 μM MitoProbe on dissociated neurons (n=3 replicates). Statistical analyses included one-way ANOVA with Dunnett’s post-hoc for time effects, two-way repeated-measures ANOVA for treatment×time interactions, and unpaired t-tests (GraphPad Prism 9; p<0.05 significance).

### 3.7. Inhibitor Studies for EV Uptake Mechanisms

To elucidate the endocytic pathways mediating uptake of Cy5-DBCO-labeled BR-EVs in primary neuronal cultures, pharmacological inhibition experiments were conducted in parallel for Cy5-DBCO-SD-EVs (from normotensive SD rats) and Cy5-DBCO-DSS-EVs (from hypertensive DSS rats). Primary neuronal cultures (10–14 days in vitro, as described previously) were pre-treated with inhibitors or vehicle control for 30 min at 37°C in phenol red-free Neurobasal medium. Dynasore (80 μM final concentration; a small-molecule inhibitor of dynamin GTPase activity, disrupting vesicle scission in clathrin- and dynamin-dependent pathways) or nystatin (50 μM final; a polyene antifungal that sequesters cholesterol, thereby disrupting caveolae formation and caveolin-mediated endocytosis) was added from 1000× DMSO stocks (final DMSO 0.1%). Vehicle controls received equivalent DMSO volumes. Cultures were then incubated with Cy5-DBCO-BR-EVs (10 μg/mL total protein) for 24 h at 37°C in 5% CO₂, followed by three PBS washes to remove unbound EVs.

Uptake was quantified by measuring mean Cy5 fluorescence intensity in neuronal cells using live-cell confocal microscopy (633 nm excitation, 650–700 nm emission; 60× objective, z-stack projections as before). ROIs were semi-automatically segmented around somata (150–300 μm² area) guided by Hoechst 33242 nuclear counterstain, with background subtraction via rolling-ball algorithm. Intensities were normalized to concurrent vehicle-treated controls within each experiment to account for batch variability, yielding % uptake reduction relative to vehicle (100% uptake). At least 15 random fields (≥50 neurons/field) were analyzed per coverslip across n=3 independent cultures per condition/EV variant/inhibitor combination.

Cell viability was assessed concurrently via 3-(4,5-dimethylthiazol-2-yl)-2,5-diphenyl-tetrazolium bromide (MTT) assay to confirm inhibitor specificity and exclude cytotoxicity. Post-incubation, MTT (0.5 mg/mL) was added for 4 h at 37°C, formazan crystals solubilized in DMSO, and absorbance read at 570 nm (reference 650 nm) on a microplate reader. Viability was expressed as % of vehicle control (n=3 replicates per condition). One-way ANOVA with Dunnett’s multiple comparisons tested inhibitor effects vs. vehicle within each EV variant (α=0.05). Two-way ANOVA assessed interactions between EV type (SD vs. DSS) and inhibitor (Dynasore vs. nystatin vs. vehicle) on % uptake reduction.

### 3.8. FRAP assay for Post-Uptake Trafficking

To evaluate the stability and diffusivity of internalized Cy5-DBCO-BR-EVs post-mitochondrial docking, FRAP experiments were conducted on live neuronal cultures at 24 h post-EV incubation (as above, without inhibitors). Selected ROIs (circular regions, 2–3 μm diameter) encompassing Cy5 puncta colocalizing with MitoTracker Green-labeled mitochondria were photobleached using a 633 nm laser at 100% power for 5 iterations (dwell time 1.27 μs/pixel). Recovery of fluorescence was monitored over 5 min with low-intensity imaging (5% laser power, 1 frame/3 s). Diffusion coefficients (D) were calculated from the initial recovery slope using the equation D = (w² / 4τ½), where w is the ROI radius and τ½ is the half-recovery time, fitted via nonlinear regression in ImageJ FRAP plugin. At least 20 ROIs per condition were analyzed across n=3 cultures per EV variant. Half-recovery times (τ½) were directly compared via unpaired t-test (p<0.05).

### 3.9. Proteomic Analysis of BR-EVs

To delineate differences in protein cargo between BR-EVs from normotensive SD and hypertensive DSS rats, untargeted liquid chromatography-tandem mass spectrometry (LC-MS/MS) was performed on purified EV samples (∼50 μL each in PBS). BR-EVs were isolated via differential ultracentrifugation as described previously. Sample preparation involved lysis in 150 μL 1× RIPA buffer with 1× Halt protease inhibitor cocktail, followed by probe sonication (3 × 10 s pulses on ice, 10 s rests). Total protein was quantified by BCA assay, with 10 μg per sample buffer-exchanged thrice into 25 mM ammonium bicarbonate (AmBic) using 50 kDa spin filters. Samples were dried, resuspended in 50 μL 25 mM AmBic/50% acetonitrile (ACN), and digested overnight at 37°C with 1 μg trypsin and 500 ng Lys-C.

Residual RIPA detergents were removed via ethyl acetate extraction: 1 mL water-saturated ethyl acetate was added per sample, vortexed (30 s), and centrifuged (14,000 × g, 15 s); the organic layer was discarded. This was repeated five times, after which the aqueous layer was dried and resuspended in 2% ACN/0.1% formic acid (FA) for LC-MS/MS.

Peptides were separated on a C18 EasySpray column (50 cm × 75 μm, 2 μm particles) using a 2-h gradient (0.1% FA in water [A] to 0.1% FA in ACN [B]): 0–3 min 3% B, 3–93 min 3–32% B, 93–103 min 32–90% B, 103–108 min 90% B, 108–120 min 3% B. Eluates were analyzed on a Q Exactive HF-X Orbitrap: full MS (m/z 350–1,600, resolution 120,000, AGC target 3e6, 20 ms max IT) followed by top-20 data-dependent MS2 (HCD, resolution 30,000, AGC 1e5, 50 ms max IT, 1.6 m/z isolation, 27% NCE).

Raw files were processed in Proteome Discoverer 2.5.0 with SEQUEST HT against the reviewed UniProt Rattus norvegicus database (UP000002494; 20,379 entries, including trypsin/Lys-C). Dynamic modifications included phosphorylation (STY; +79.9663 Da); static: carbamidomethyl (C; +57.0215 Da). Precursor/ fragment mass tolerances: 10 ppm/0.02 Da. Percolator FDR was set at 1% (protein/PSM/peptide). Proteins were grouped by parsimony; shared/unique assignments based on sample-specific detection. Gene Ontology (GO) terms, Pfam domains, and pathways (Reactome/WikiPathways) were annotated via integrated nodes. Phosphosites were localized with ptmRS (≥75% confidence). No label-free quantification was applied; analysis focused on qualitative identification and GO/pathway enrichment (term frequency via semicolon-split parsing).

### 3.10. ICV Injection of Cy5-DBCO-SD-EVs

Unilateral injections targeted the right lateral ventricle using stereotaxic guidance (Kopf Instruments). SD Rats were induced with 5% isoflurane in O₂ and maintained at 2.5% during the procedure, with body temperature stabilized at 37°C via a heated platform. The scalp was incised along the midline, and the periosteum cleared to expose the skull. Bregma and lambda were zeroed for alignment, followed by a burr hole at AP: −0.8 mm, ML: +1.6 mm, DV: −3.6 mm. A 10-μL Hamilton syringe (33-gauge needle) delivered 10 μg Cy5-DBCO-SD-EVs (in PBS) at 1 μL/min via an UltraMicroPump 3 (World Precision Instruments), with a 5-min post-infusion dwell. Wounds were closed with sutures, and animals recovered under observation.

At 24 h post-injection, rats were terminally anesthetized (ketamine/xylazine, 100/10 mg/kg i.p.) and transcardially perfused with ice-cold PBS (200 mL) then 4% PFA (300 mL). Brains were dissected, post-fixed in 4% PFA (overnight, 4°C), equilibrated in 30% sucrose/PBS, embedded in O.C.T. (Sakura Finetek), and cryosectioned (Leica CM1950) at 15 μm for coronal slices encompassing cortex and hippocampus.

### 3.11. Immunofluorescence Staining and Imaging

Sections were processed with minor adaptations from prior protocols. ^37^After PBS rinses (3 × 10 min), blocking proceeded in 5% horse serum/PBS (30 min, RT). Primary incubation used rabbit anti-NeuN (1:300; Cell Signaling), anti-GFAP (1:500; Abcam), or anti-IBA1 (1:500; Fujifilm Wako) diluted in PBS + 0.5% Triton X-100 + 5% horse serum (48 h, 4°C). Post-rinse (PBS, 3 × 10 min), slides received Alexa Fluor 488 donkey anti-rabbit IgG (1:500; Thermo Fisher A-21206; 1 h, RT). DAPI (1:5000) counterstained nuclei.

Slides were mounted and imaged via confocal microscopy. Z-stacks (1-μm steps) captured NeuN/GFAP/IBA1 immunoreactivity and Cy5 signal for EVs tracking. Quantitative fluorescence analysis followed ImageJ protocols, with ROIs defined anatomically and intensities normalized to area after background subtraction.

### 3.12. Synthesis of Cy5-DBCO

#### 3.12.1. Synthesis of Compound 1: N-((1E,3E)-3-(phenylimino)prop-1-en-1-yl)aniline (Malonaldehyde Dianil Precursor)

To a stirring solution of distilled water (20 mL), HCl (4 mL), and 1,1,3,3-tetramethoxypropane (2.0 g, 0.022 mol) was added dropwise a mixture of distilled water (20 mL), HCl (5 mL), and aniline (3.0 g, 0.020 mol), followed by continued stirring at ∼50 °C for 3 h. The reaction mixture was filtered, and the filtrate was washed with ether to afford compound 1 (1.76 g, 50%) as an orange powder. ¹H NMR (400 MHz, DMSO-d₆) δ 6.53 (t, J = 11.5 Hz, 1H), 7.16 (t, J = 7.2 Hz, 2H), 7.26–7.62 (m, 8H). ¹³C NMR (101 MHz, DMSO-d₆) δ 159.00, 139.31, 130.42, 126.44, 118.04, 99.34.

#### 3.12.2. Synthesis of Compound 2: 2,3,3-Trimethyl-3H-indole

Compound 2 was prepared through a classic Fischer indole synthesis by reacting phenylhydrazine with 3-methylbutanone (isopropyl methyl ketone) in glacial acetic acid. The mechanism begins with the formation of phenylhydrazine acetate, which facilitates imine formation with the ketone. Subsequent acid-catalyzed protonation of the imine leads to a [3,3]-sigmatropic rearrangement, yielding an enamine intermediate that undergoes intramolecular electrophilic aromatic substitution and dehydration to furnish the indole core. In brief, phenylhydrazine (1.62 g, 0.015 mol) was dissolved in glacial acetic acid (25 mL), and 3-methylbutanone (2.5 mL, 0.017 mol) was added to the stirring solution. The mixture was refluxed for several hours until completion (monitored by TLC). The reaction was then cooled, and the solvent was removed by rotary evaporation under reduced pressure. The residue was partitioned between diethyl ether (30mL) and water (30 mL), with the aqueous layer re-extracted with additional ether (30 mL). The combined organic layers were dried over anhydrous sodium sulfate, filtered, and concentrated under vacuum to yield a brownish liquid. This crude product was further purified by short-path distillation if necessary, providing Compound 2 (4.0 g, 83% yield). ¹H NMR (400 MHz, CDCl₃) δ 1.26 (s, 6H), 2.24 (s, 3H), 7.15 (t, J = 7.4 Hz, 1H), 7.25 (dd, J = 8.9, 4.1 Hz, 2H), 7.51 (d, J = 7.6 Hz, 1H). ¹³C NMR (101 MHz, CDCl₃) δ 188.17, 153.56, 145.69, 127.74, 125.30, 121.45, 119.99, 119.68, 53.84, 23.37, 15.62.

#### 3.12.3. Synthesis of Compound 3: 1-Ethyl-2,3,3-trimethyl-3H-indol-1-ium Iodide

Compound 3, a quaternized indoleninium salt, was generated from Compound 2 via an SN2 alkylation with ethyl iodide in acetonitrile. The pyrrole nitrogen of the indole, which is deprotonated under the reaction conditions, acts as a nucleophile, displacing iodide from the ethyl electrophile to form a new C–N bond. The resulting quaternary ammonium stabilizes through delocalization into the indole ring, with no further rearrangement required. Procedurally, Compound 2 (2.0 g, 12.5 mmol) was dissolved in dry acetonitrile (25 mL) under a nitrogen atmosphere. Ethyl iodide (2.5 mL, ∼20 mmol) was added dropwise, and the mixture was refluxed at ∼82 °C for 48 hours. Progress was monitored by TLC (silica gel, ethyl acetate/hexane). After cooling to room temperature, the solvent was removed by rotary evaporation, and the orange oily residue was triturated with diethyl ether (3 × 25 mL) to precipitate the product. The solid was collected by filtration, washed with cold ether, and dried under high vacuum, yielding compound 3 (85% yield) as an orange powder suitable for direct use in subsequent steps. ¹H NMR (400 MHz, CDCl₃) δ 1.54–1.58 (m, 9H; overlapping methyl singlets and ethyl methylene), 3.06 (s, 3H; 2-methyl), 4.66 (q, J = 7.3 Hz, 2H; N-CH₂), 7.51 (m, 3H; aromatic), 7.54 (m, 1H).

#### 3.12.4. Synthesis of Compound 4: 1-(5-Carboxypentyl)-2,3,3-trimethyl-3H-indol-1-ium Inner Salt

The preparation of Compound 4, a carboxylated indoleninium salt, proceeded in a two-step, one-pot manner starting from 6-bromohexanoic acid. First, a Finkelstein halide exchange with potassium iodide generates 6-iodohexanoic acid as the alkylating agent (noted as an “acyl iodide” in some literature but functioning as an ω-iodoalkyl carboxylic acid). This intermediate then undergoes nucleophilic addition to the enamine tautomer of 2,3,3-trimethyl-3H-indole, forming a tetrahedral adduct that rearranges to a zwitterionic iminium-carboxylate species. Proton loss from the α-position yields the final indolium structure. In brief, potassium iodide (1.85 g, 11 mmol) was dissolved in dry acetonitrile (20 mL), and 6-bromohexanoic acid (1.0 g, 5 mmol) was added dropwise with stirring. The mixture was heated to 50 °C for 30 minutes to facilitate the exchange. Subsequently, 2,3,3-trimethyl-3H-indole (0.8 g, 5 mmol) was introduced, and the reaction was refluxed overnight under nitrogen. After cooling to room temperature, the mixture was filtered to remove KBr, and the filtrate was concentrated by rotary evaporation. The residue was partitioned between dichloromethane (25 mL) and water (25 mL); the organic layer was dried over sodium sulfate and evaporated. The crude product was then precipitated from ether, filtered, and dried, affording Compound 4 (75% yield) as a gray solid. ¹H NMR (400 MHz, DMSO-d₆) δ 1.37 (d, J = 17.5 Hz, 2H; apparent due to overlap), 1.51 (m, 6H; gem-dimethyl and chain CH₂), 1.82 (p, J = 7.9 Hz, 2H; chain CH₂), 2.19 (t, J = 7.4 Hz, 2H; CH₂COOH), 2.47 (s, 3H; 2-methyl), 4.43 (t, J = 7.7 Hz, 2H; N-CH₂), 7.56–7.64 (m, 2H; aromatic), 7.73–7.89 (m, 1H; aromatic), 8.00-8.90 (m, 1H). ¹³C NMR data aligns with the zwitterionic form.

#### 3.12.5. Synthesis of Asymmetric Cyanine Dye 5

Asymmetric cyanine dye 5, featuring a pentamethine bridge and delocalized chromophore, was assembled via a stepwise condensation of the unsymmetrical indoleninium salts (compounds 3 and 4) with malonaldehyde dianil (compound 1). The process integrates the quaternary ammonium from compound 3 into the polymethine chain through imine formation and dehydration, while the carboxypentyl chain from compound 4 remains pendant. A final alkylation step quaternizes any residual tertiary amine on the bridge. In brief, compound 3 (1.0 g, 3.2 mmol) was suspended in a mixture of glacial acetic acid (10 mL) and acetic anhydride (5 mL), and Compound 1 (1.0 g, 3.8 mmol) was added portionwise under nitrogen. The mixture was refluxed at ∼120 °C for 1 hour, cooled to room temperature, and then treated with a solution of Compound 4 (2.0 g, 5.0 mmol) in anhydrous pyridine (20 mL). Stirring continued at room temperature overnight under nitrogen. The deep green solution was poured into cold diethyl ether (100 mL) to precipitate the crude product, which was collected by filtration, washed with ether, and dried under vacuum. The green solid (∼3.2 mmol, ∼100% crude yield based on limiting reagent) was used directly without further purification due to its stability and purity sufficient for conjugation. The crude dye exhibits characteristic absorption at ∼650 nm (ε ≈ 2.5 × 10⁵ M⁻¹ cm⁻¹ in MeOH) and emission at ∼670 nm, confirming the Cy5-like structure.

#### 3.12.6. Synthesis of Cy5-DBCO

The carboxylic acid functionality of compound 5 was activated for amide coupling with Dibenzocyclooctyne-amine (DBCO-NH₂), a strained alkyne derivative for copper-free click chemistry, yielding Cy5-DBCO as a mitochondrial-targeted fluorescent probe. This step mirrors standard peptide coupling protocols, forming a stable amide linkage while preserving the dye’s chromophore and the DBCO’s reactivity. Adapting established conditions, crude Compound 5 (∼3.2 mmol) was dissolved in dry dichloromethane (50 mL) or DMF (50 mL) under nitrogen. DBCO-amine (1.2 equiv., ∼3.8 mmol, typically 1.0 g) was added, followed by a coupling agent such as HBTU (2.0 g, 5.0 mmol) or EDC/NHS (1.2 equiv. each), and triethylamine (5 mL, base) or DIPEA. The mixture was stirred at room temperature overnight, monitoring by TLC (DCM/MeOH 20:1). The solvent was removed by rotary evaporation, and the residue was purified by flash column chromatography on silica gel (eluent: DCM/MeOH gradient from 40:1 to 20:1) or preparative HPLC (C18, MeOH/H₂O with 0.1% TFA). Fractions containing the product (UV-active at 650 nm) were combined, evaporated, and lyophilized to afford Cy5-DBCO as a green solid (yield: 70–85%, depending on purity of 5). Cy5-DBCO has been validated as a mitochondrial-targeted dye, showing high colocalization with MitoTracker in vitro (HeLa cells).

1H NMR (500 MHz, CDCl3) δ ppm 1.07-1.24(m, 1H),1.33-1.45(m, 5H), 1.54-1.63(m, 2H), 1.63-1.83(m, 14H), 1.91-2.08(m, 3H), 2.43-2.51 (m, 1H), 3.17-3.39 (m, 2H), 3.60-3.68(m, 1H), 3.87-4.01(m, 2H), 4.01-4.12(m, 2H), 5.06-5.14(m, 1H), 6.05-6.20(m, 3H) 6.55-6.63 (m, 1H), 7.01-7.14(m, 2H), 7.14-7.26(m, 3H), 7.27-7.42(m, 9H), 7.61-7.68(m, 1H), 7.89-8.00 (m, 2H):13C NMR (101 MHz, DMSO-d6) δ 172.92 172.76 172.57 171.94 165.73 153.49 153.29 151.07 148.04 141.93 141.54 141.43 141.27 132.14 129.17 128.79 128.65 128.62 128.37 128.26 127.82 127.09 125.80 125.47 125.26 125.17 122.94 122.35 122.31 122.27 114.70 110.51 110.41 107.91 103.20 103.13 77.45 77.39 77.19 76.93 55.45 53.56 49.46 49.38 43.99 39.18 38.59 36.02 35.26 34.68 27.84 27.73 26.99 26.32 24.99 12.26.

## 4. Conclusion

This study establishes SPAAC as a biocompatible, high-fidelity chemical platform for engineering mitochondria-targeted BR-EVs without perturbing their structural, biochemical, or functional integrity. Through low-density Cy5-DBCO conjugation (10–20% surface occupancy; 72–75% recovery), SPAAC enabled site-specific functionalization under copper-free, aqueous conditions, preserving nanoscale morphology (TEM: 30–150 nm; DLS: 120–200 nm; p > 0.05) and eliminating oxidative or aggregation artifacts.

Complementary time-course experiments demonstrated unaltered uptake kinetics and efficient intracellular routing. SPAAC-engineered BR-EVs internalized predominantly via bulk endocytosis and dynamin-dependent macropinocytosis,^31^ exhibiting linear accumulation over 48 hours and >75% mitochondrial colocalization (Pearson’s r = 0.82). The FRAP assay confirmed maintained vesicle mobility upon mitochondrial docking, indicating preserved membrane dynamics and fusion competence. Importantly, SPAAC labeling did not affect organelle morphology—SD-EVs maintained elongated mitochondrial networks (aspect ratio 4.0–4.3), whereas DSS-EVs induced fragmentation (aspect ratio 2.1; p < 0.001), reflecting intrinsic cargo differences rather than chemical modification.

Oxidative stress assays further revealed that DSS-derived EVs elevated mitochondrial reactive oxygen species through NOX2/CYBA activation (1.6-fold increase; p < 0.05),^38^ while SD-EVs sustained redox balance—effects independent of SPAAC labeling. Proteomic profiling of unlabeled SD- and DSS-EVs (760 total proteins, FDR < 1%) confirmed this functional divergence, identifying 145 DSS-unique proteins enriched in oxidative and complement pathways (e.g., HSPA1L, GSTM7, C3, Rab1B/8B/10) compared with SD-EVs enriched in cytoskeletal and metabolic maintenance factors.

In vivo analyses reinforced SPAAC’s biocompatibility and translational safety. Neuroinflammation assays revealed no increases in GFAP, IBA1, or CD68 immunoreactivity following intracerebroventricular administration of Cy5-DBCO–SD-EVs, confirming the immunological neutrality of the modification. Biodistribution imaging showed strong CNS retention and minimal peripheral dispersion 24 hours post-injection, consistent with glymphatic-mediated clearance rather than reticuloendothelial sequestration.

These findings demonstrate that SPAAC modification preserves exosomal architecture, trafficking dynamics, and biological identity while enabling precise, mitochondria-targeted labeling. The integration of proteomics, live-cell imaging, FRAP mobility analyses, and in vivo validation positions SPAAC as a chemically orthogonal and biologically inert framework for functionalizing exosomes with organelle-specific precision. This approach establishes a foundational platform for mechanistic visualization and future therapeutic development targeting mitochondrial dysfunction in hypertension and other neuroinflammatory disorders.

## 5. Limitations

Our present study provides compelling validation of SPAAC as a biocompatible and high-fidelity strategy for functionalizing BR-EVs toward mitochondrial targeting. While the results demonstrate excellent preservation of vesicle integrity, bioactivity, and in vivo compatibility, several aspects remain to be further developed as the platform moves toward broader application.

First, the current work was designed to establish mechanistic fidelity and biological safety, rather than to model full systemic pharmacokinetics. The use of neuronal and neuron–astrocyte co-culture systems, together with intracerebroventricular administration, allowed controlled evaluation of cellular uptake, mitochondrial interaction, and biocompatibility in the absence of systemic confounders. Future studies incorporating intravenous or targeted delivery paradigms will extend these findings to clinically relevant routes.

Second, Cy5-DBCO–based fluorescence imaging effectively captured uptake kinetics, mitochondrial docking, and biodistribution. However, expanding to multi-color or multimodal labeling could further enrich spatial and temporal resolution of exosomal trafficking and cargo turnover in complex tissue environments.

Third, label-free proteomic profiling successfully delineated the molecular divergence between normotensive and hypertensive BR-EVs, identifying oxidative and complement pathway enrichment in DSS-derived vesicles. Incorporating quantitative and post-translational proteomic workflows in future studies will refine understanding of regulatory modifications underlying disease-specific signaling.

Finally, this study focused on SPAAC conjugation using a single mitochondrial-targeting fluorophore as a model ligand. The chemistry’s modularity now opens opportunities for conjugating bioactive peptides, redox sensors, or therapeutic cargos to create multifunctional exosome systems for precision mitochondrial modulation. These planned extensions represent natural progressions of the platform, rather than experimental constraints, and will further define SPAAC-engineered exosomes as a scalable and translationally adaptable nanotechnology for organelle-targeted delivery.

## ASSOCIATED CONTENT

## Supporting Information

## Author Contributions

The manuscript was written through contributions of all authors. All authors have given approval to the final version of the manuscript. ‡These authors contributed equally.

## ACKNOWLEDGMENT

We sincerely thank the NIH (R01HL163159, Z.S.), NIH (R15 EB035866, L.B), and the American Heart Association (AHA, grant #1807047, LB), GLRC-ICC (L.B., R01805) for their generous financial support. We also extend our great gratitude to Dr. Rick Koubek for his encouragement and unwavering support throughout our project.

